# The visual white matter connecting human area prostriata and the thalamus is retinotopically organized

**DOI:** 10.1101/2020.04.19.047993

**Authors:** Jan W. Kurzawski, Kyriaki Mikellidou, Maria Concetta Morrone, Franco Pestilli

**Affiliations:** IRCCS Stella Maris, Calambrone, Pisa, Italy; Department of Psychology & Center for Applied Neuroscience, University of Cyprus, Cyprus; Department of Translational Research on New Technologies in Medicine and Surgery, University of Pisa, Italy; Department of Psychological and Brain Sciences, Program in Neuroscience and Program in Cognitive Science, Indiana University, 1101 E 10th Street, Bloomington, IN 47401

## Abstract

The human visual system is capable of processing visual information from fovea to the far peripheral visual field. Recent fMRI studies have shown a full and detailed retinotopic map in area prostriata, located ventro-dorsally and anterior to the calcarine sulcus along the parietooccipital sulcus with strong preference for peripheral and wide-field stimulation. Here, we report the anatomical pattern of white-matter connections between area prostriata and the thalamus encompassing the lateral geniculate nucleus (LGN). We observe a continuous and extended bundle of white matter fibers from which two subcomponents can be extracted: one passing ventrally parallel to the optic radiations (OR) and another passing dorsally circumventing the lateral ventricle. Interestingly, the loop travelling dorsally connects the thalamus with the central visual field representation of prostriata, while the other loop travelling more ventrally connects the LGN with the more peripheral visual field representation. This is consistent with a retinotopic segregation recently demonstrated in the OR, connecting the LGN and V1 in humans. Our results demonstrate for the first time a retinotopic segregation regarding the trajectory of a fiber bundle between the thalamus and an associative visual area.

## Introduction

Mikellidou, Kurzawski et al. (2017b) described for the first time the functional properties of area prostriata in humans using a novel wide-field visual stimulation system (Greco et al. 2016). In humans, area prostriata is located at the junction between the calcarine and parietooccipital (POS) sulci, adjacent to the far peripheral representation of V1 and ventral V2 (Mikellidou et al. 2017b). Area prostriata preferentially responds to very fast motion, greater than 500 deg/sec, and has a complete representation of the visual field, clearly distinct from the adjacent area V1. Interestingly, area prostriata shows little, if any, cortical magnification but rather has a homogeneous distribution of receptive field centers across eccentricity. The posterior-dorsal border of human prostriata—abutting V1—represents the far peripheral visual field, with eccentricities decreasing toward its anterior boundary in the fundus of the calcarine sulcus. This is an anatomical organization comparable to that reported in the common marmoset (Yu et al. 2012). The functional properties of area prostriata suggest that the area may serve to alert the brain about fast visual events, particularly in the peripheral visual field and it could subserve as a critical node for transient attention (Liu et al. 2005).

The cortical position, structural and functional properties of human area prostriata (Ding et al. 2016; Glasser et al. 2016; Sanides 1969) are similar to other species (Kobayashi and Amaral 2003; Morecraft et al. 2000; Rosa et al. 1997; Sanides and Vitzthum 1965). In humans, area prostriata shows lower myelination and increased cortical thickness as compared to neighbouring visual areas (see, Figure 4F-G in Glasser et al. (2016)). The area shows characteristics akin to those of the limbic cortex (Rockland 2012), as it lacks the clearly defined laminar profile evident throughout the cerebral cortex, with a thinner layer 4 and a thicker layer 2 (Ding et al. 2003). Figure 1 shows coronal and sagittal planes of human area prostriata manually labelled in red on an atlas (1^st^ column) and a histological slice (2^nd^ column). A T1-weighted slice is also shown with area prostriata in white mapped from a multimodal atlas (3^rd^ column).

**Figure 1.**
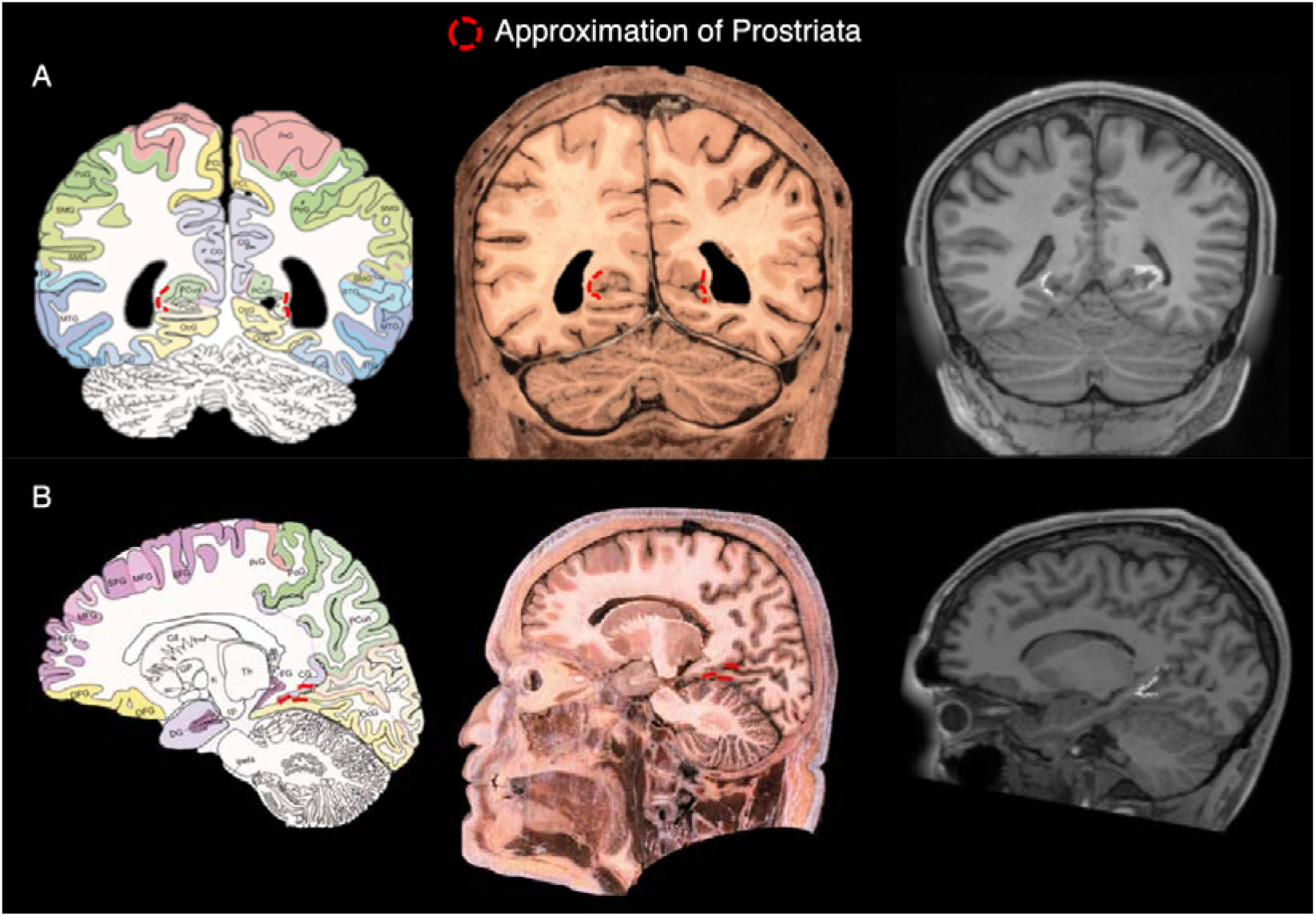
Putative area prostriata labelled in red trace on a labelled atlas, a histological slice (Mai 2016) and a T1-weighted slice from one of our subjects. In T1-weighted image Prostriata has been mapped from a multi-modal parcellation template (Glasser et al. 2016). A. Coronal view. B. Sagittal view.

As of today, little is known about the anatomical connections of human area prostriata. The connections of area prostriata have been reported in animal models using retrograde and anterograde tracers but to date, these are unknown in the human brain. Prostriata connections, including the motion-sensitive MT+, auditory cortices, cingulate motor areas, the orbitofrontal cortex, and the frontal polar cortices (Burman et al. 2011; Falchier et al. 2010; Morecraft et al. 2000; Palmer and Rosa 2006), support the proposal that it can act as an interface for the coordination of motor responses required in orienting, and could mediate attentional shifts, postural and defensive reactions in ‘fight’ or ‘flight’ situations. In humans, it is challenging to establish whether prostriata has a connection (input and/or output) to the thalamus, using the available non-invasive techniques. Recently published work that examined connections of prostriata in rodents showed more evident connections between peripheral V1 and prostriata in comparison to lateral, more foveal V1 (Lu et al. 2019). Moreover, Lu et al., proposes that prostriata might be directly connected to the thalamus, as previously suggested (Ding 2013).

Our aim here is to use computational tractography to explore the structural connectivity of human area prostriata with a portion of the thalamus that includes the lateral geniculate nucleus (LGN). In primates and cats, the LGN provides direct input to primary and associative visual cortex, but it has also been shown to receive numerous feedback projections from afferent cortical fibers of the visual cortex (Florence and Casagrande 1987; Sherman and Guillery 2002). To investigate the structural connectivity of area prostriata with the LGN, we combined three different data modalities (T1-weighted, T2-weighted, and functional MRI) with the use of modern tractography, statistical evaluation methods and Human Connectome Project (HCP) data. We performed tractography, evaluated the results using LiFE (Pestilli et al. 2014) and then combined the functionally defined maps, the myelin content and cortical thickness maps together with tractography to fully characterize the corticothalamic connections of area prostriata.

## Results

### Estimating eccentricity, myelin-content and cortical thickness of area prostriata using public datasets and atlases

We set to report the relationship between functional and anatomical properties of area prostriata using published datasets and atlases (Mikellidou et al. 2017b; Benson et al. 2018; Glasser et al. 2016). We focused on eccentricity maps (Mikellidou et al. 2017b; Benson et al. 2018), myelin-sensitive maps (Hagiwara et al. 2018; Shams et al. 2019; Glasser and Van Essen 2011) and cortical thickness maps (Glasser et al. 2016). The retinotopic organization of area prostriata estimated using functional neuroimaging shows a typical pattern (Mikellidou et al. 2017b; Yu et al. 2012). It has been reported that the visual field periphery is located in the most posterior region of prostriata, abutting V1 periphery (Figure 2A). The central portion of the visual field is located in the anterior portion of prostriata. This indicates the presence of a posterior to anterior eccentricity gradient in area prostriata. This reversal is a canonical property of visual field organization in human visual cortex (Wandell and Winawer 2015; Wandell et al. 2005).

We investigated the degree of generality of prostriata’s anterior-posterior eccentricity axis using an independent dataset (Glasser et al. 2016). The human brain atlas published by Glasser et al. (2016) contains a map of prostriata (ProS ROI). We show that the location of prostriata estimated by Mikellidou, Kurzawski et al. (2017b) is very similar to the one reported by Glasser and colleagues (2016). To do so, we report the barycentre of prostriata in Talairach coordinates as provided by Glasser et al., (2016): X: −22.0, Y: −55.7, Z: 2, and compare these coordinates to the barycentre estimated by Mikellidou, Kurzawski et al. (2017b) X: −19.8 ± 1.7, Y: −60.7 ± 3.0, Z: 1.4 ± 4.5. The Talairach coordinates show a high degree of overlap. This is noticeable especially given the differences in mapping method and resolution used in the two studies.

We next investigated whether an anterior-posterior axis can be observed within area prostriata from eccentricity, myelin-sensitive and cortical thickness maps (Figure 2B-D) using data from the Human Connectome Project (Glasser et al. 2016; Benson et al. 2018). Anterior-posterior axes (map gradients) are visible in each map. First, we replicated the existence of an anterior-posterior eccentricity gradient in the HCP dataset (Figure 2B). In addition to that, we also show that a similar anterior-posterior axis exists in the myelin-sensitive maps (Figure 2C) and in the cortical thickness maps (Figure 2D). In sum, the most anterior part of prostriata processes the central visual field, contains less myelin and has higher cortical thickness, while the posterior part of prostriata processes the peripheral visual field, has higher myelin content and lower cortical thickness. A similar orderly relationship between eccentricity, cortical thickness and myelin content maps has also been previously demonstrated for other visual areas (Abdollahi et al. 2014; Burge et al. 2016). These published data provide an underdescribed anterior-posterior axis in area prostriata consistent across multiple data modalities.

**Figure 2.**
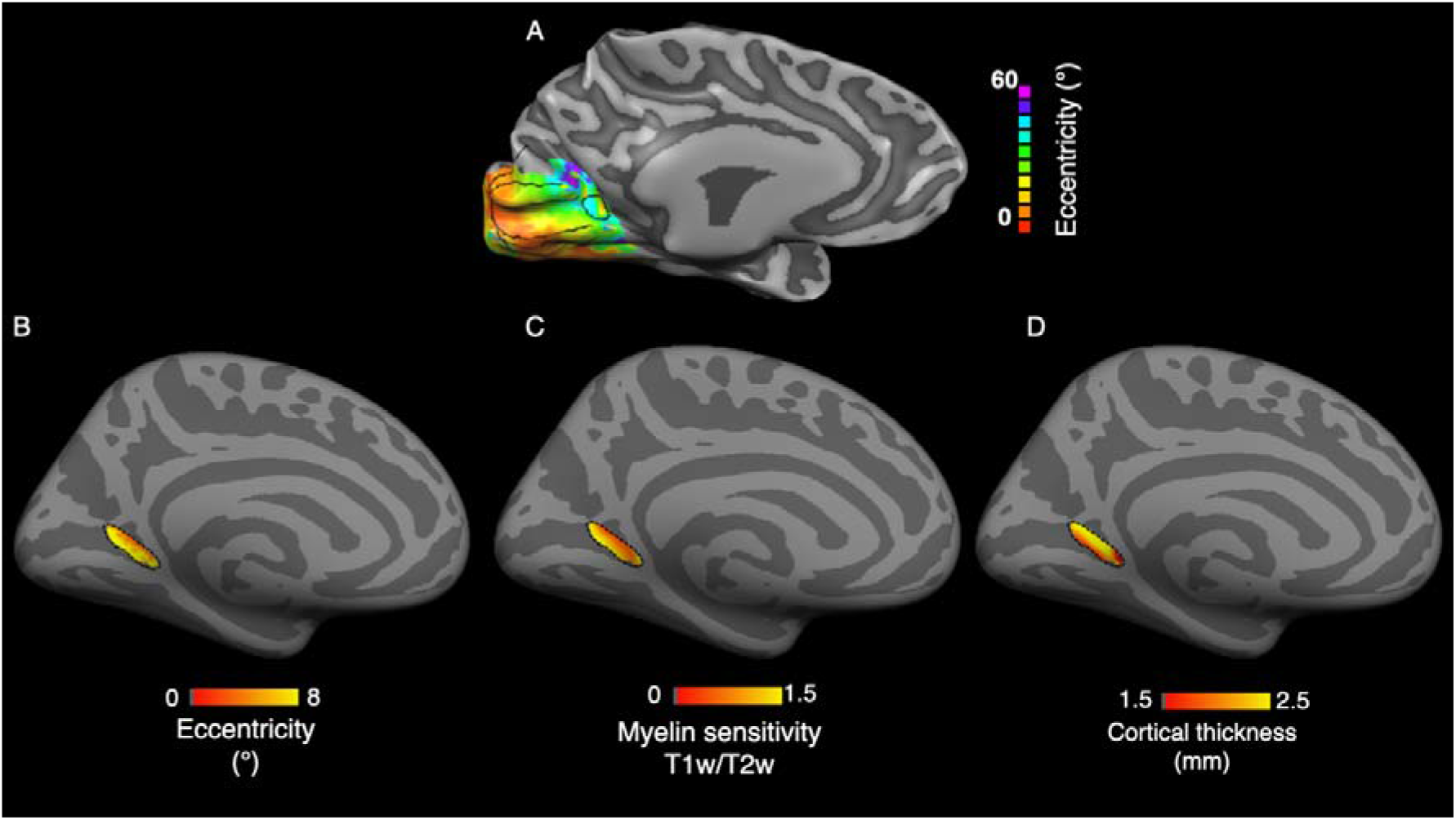
Structural and functional properties of area prostriata. A. Eccentricity representation of subject (S8) from Mikellidou et al., (2017b) with visual field stimulation extending up to 60 degrees. Black lines represent the division between visual areas with a distinct area prostriata located at the fundus of calcarine sulcus. B. Eccentricity representation (N=181) within area prostriata with a visual field stimulation extending up to 8 degrees using HCP data (Benson et al. 2018). C. Myelin content and D. cortical thickness maps (N=210) within area prostriata created using HCP data (Glasser et al. 2016)

### Visual white matter connections between prostriata and the thalamus

In this section, we set to identify the visual white matter connections between area prostriata and the thalamus. We used multishell DWI data (Glasser et al. 2013; Andersson et al. 2003; Andersson and Sotiropoulos 2015; Andersson and Sotiropoulos 2016) and ensemble tractography (Caiafa and Pestilli 2017; Pestilli 2015; Pestilli et al. 2014; Takemura et al. 2016a; Smith et al. 2012) to track between a subcortical region comprising the LGN and prostriata (see Methods for additional details). Figure 3A shows the white matter tract connecting the LGN and prostriata. Tracking results show a large and anatomically complex tract that we refer to as LGN-Pro (Figure 3A in green).

We note that the tract has a complex anatomical trajectory, with fibers travelling both superiorly and laterally. We also notice that anatomical trajectory of the fibers within the LGN-Pro tract resembles (but only partially overlaps with) that of the optic radiations (OR). It has been shown that the OR can be subdivided into at least two sub-components defined by the termination of the fibers into V1 regions representing different eccentricities (Parraga et al. 2012; Yoshimine et al. 2018). The fibers of the OR projecting to V1 locations representing the visual periphery travel more ventrally and laterally within Meyer’s loop. The fibers of the OR projecting to V1 locations representing the central visual field travel more dorsally and medially outside of Meyer’s loop (Yoshimine et al. 2018). Here, we used the anterior-posterior axis of prostriata described in the previous section (Figure 2) and subdivided the LGN-Pro tract into anterior projecting and posterior projecting fibers (Figure 3B yellow and cyan respectively). We call the component projecting to anterior prostriata LGN-Pro-1 (yellow) and the component projecting to posterior prostriata LGN-Pro-2 (cyan). These subcomponents were subdivided automatically in 139 subjects using their termination zone in prostriata as a criterion. LGN-Pro fiber was assigned to LGN-Pro-1 or LGN-Pro-2 depending on whether it terminated into a voxel within the anterior 50% of the prostriata voxels or the posterior 50% respectively (Figure 4). This analysis was performed automatically using brainlife.io (Avesani et al. 2019). To do this we developed an App (https://doi.org/10.25663/brainlife.app.282) that can be publicly accessed and used on new data (see Methods for additional details on the full set of Apps on brainlife.io that were used for this analysis).

A density map reports the spatial position of LGN-Pro as a whole, LGN-Pro-1 and LGN-Pro-2 across 139 (11 excluded from 150 batch) subjects aligned to the MNI template (Figure 3). Figure 3A shows that LGN-Pro is an anatomically continuous bundle across the majority of subjects. Figure 3B compares the spatial position of LGN-Pro-1 (yellow) and LGN-Pro-2 (cyan). Thresholding the map, by showing only the fibers consistently present in at least 90 subjects (65%), shows that LGN-Pro-1 (yellow) has a dorsal trajectory and LGN-Pro-2 (cyan) has a ventral trajectory. Figure 3C shows substantial overlap between LGN-Pro-2 and the OR. The exact and detailed separation of LGN-Pro-2 and OR would require other, probably histological, methods. Although the white matter bundle is shared between the two, LGN-Pro-2 originates from prostriata that is clearly different from V1 and therefore we consider it as a separate entity.

**Figure 3.**
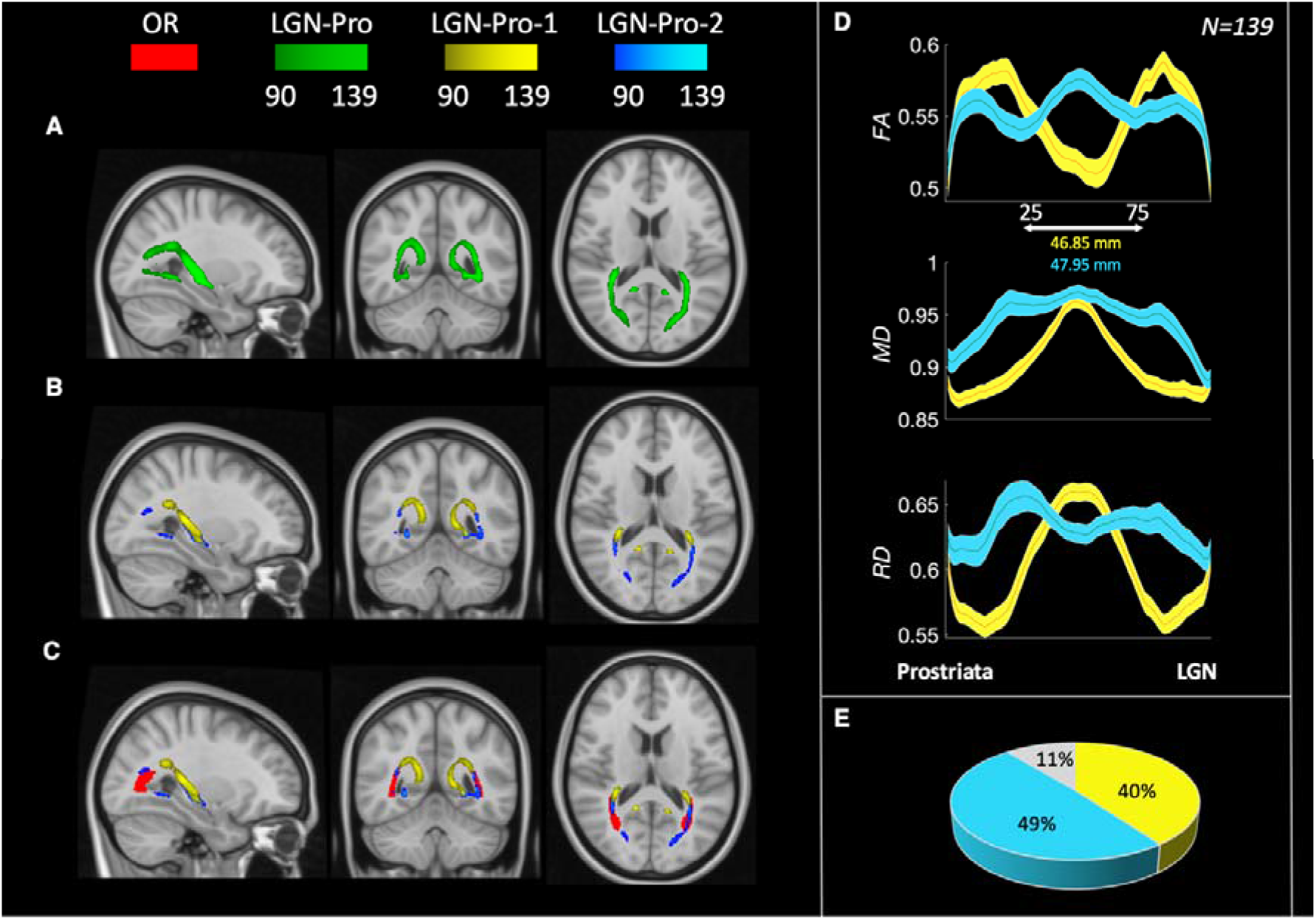
A. Map of the LGN-Prostriata tract averaged across subjects in the MNI space. B. LGN-Pro1 (yellow) and LGN-Pro2 (cyan) subcomponents averaged across subjects in the MNI space. C. LGN-Pro subcomponents shown alongside the OR (red). The colormap in A, B, C represents consistency across 139 subjects (See “Group averages” in Methods for details). D. Diffusion properties along the tract for both subcomponents. E. Percentage of total fibers between LGN and Prostriata classified as LGN-Pro-1 (yellow) and LGN-Pro-2 (cyan). In grey we report the percentage of fibers not univocally assigned to either the anterior (foveal, low-myelin, high-cortical thickness) or posterior (peripheral, high-myelin, low-cortical thickness) prostriata regions.

Finally, we used the diffusion-tensor model (Basser et al. 1994; Pierpaoli et al. 1996) and estimated fractional anisotropy (FA), mean diffusivity (MD) and radial diffusivity (RD) at 100 locations along the tracts’ length using tract profiles (Rykhlevskaia et al. 2009; Yeatman et al. 2012; Yoshimine et al. 2018). The two tracts differ substantially along their length in their microstructural properties (FA, MD, RD; Figure 3D), while the automatic subdivision of subcomponents reveals a similar number of fibers assigned to each of them (Figure 3E).

### The LGN-Pro subcomponents defined using tract trajectory, map onto prostriata locations with different properties of eccentricity, myelin-content and thickness

In the previous section we characterized the anatomy and microstructure of the complex white matter tract connecting area prostriata and the visual thalamus. We subdivided the tract into two subcomponents based on their cortical locus within prostriata (anterior/posterior). The analysis revealed two subcomponents with different anatomical trajectories and microstructural properties. Here, we investigate whether these two subcomponents show any relationship with eccentricity, myelin and cortical thickness gradient maps within prostriata (Figure 2), defined using their anatomical trajectories.

To study the relationship between LGN-Pro-1, LGN-Pro-2, and prostriata gradients (eccentricity, myelin-sensitivity and thickness) we extracted two subcomponents using a different method outside brainlife.io on an additional cohort of HCP subjects (*N=10*). Learning from the anatomical organization of the LGN-Pro tract shown in Figure 3B-C we extracted LGN-Pro-1 and LGN-Pro-2 using a manual “plane extraction” method (Zhang et al. 2010; Catani and Thiebaut de Schotten 2008). Specifically, to extract the dorsal subcomponent (LGN-Pro-1), we drew several exclusion planes (available as ROIs) to remove all fibers that travel ventrally. Similarly, to extract LGN-Pro-2 (ventral subcomponent) several exclusion planes were drawn to remove all fibers that travel dorsally (see Methods for additional details).

**Figure 4.**
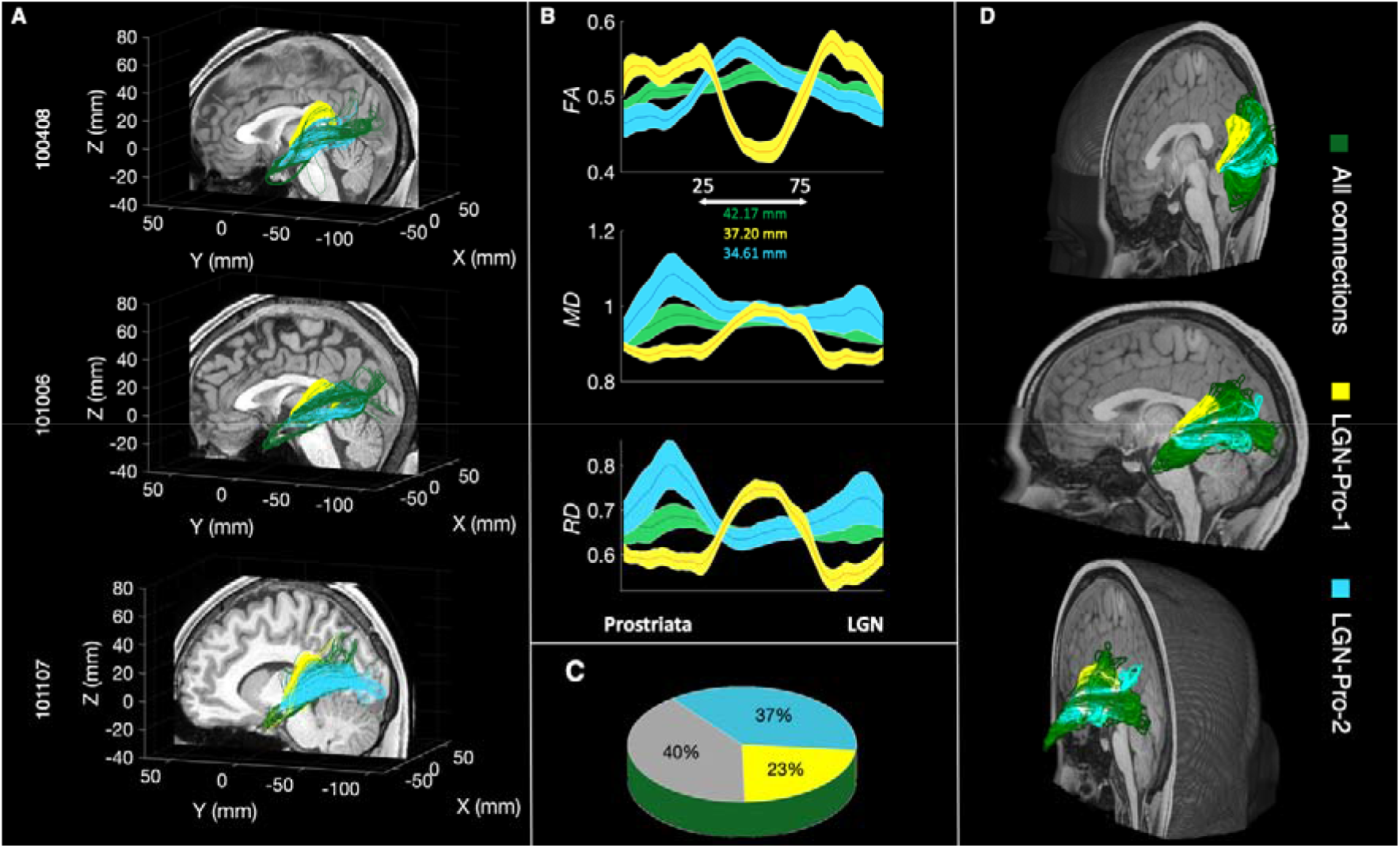
A. Shows a streamline-based representation of the white matter tract between the LGN and prostriata for three example subjects from the HCP dataset (100408, 101006, 100206). One of the two subcomponents of the tract (LGN-Pro-1 in yellow) encompasses the lateral ventricle, reaching the LGN from a superior direction. The second subcomponent (LGN-Pro-2 in cyan) follows a route similar to the OR. B. Average diffusion properties (FA, MD, RD) resampled to one hundred points for the tracts displayed in yellow and cyan. The distance between 25^th^ and 75^th^ point is 37.20 mm and 34.61 mm for LGN-Pro-1 and LGN-Pro-2 respectively (N=10). C. Proportion of fibers assigned to LGN-Pro-1 (Yellow) and LGN-Pro-2 (Cyan) using a manually curated segmentation method (plane extraction). Across subjects, the two subcomponents make up 60% of the LGN-Pro connection, with the remaining 40% intermingled between the two (N=10). D. 3D rendering for the LGN-Prostriata connection for a single subject (101309) viewed from different angles.

Figure 4A shows these two subcomponents of the LGN-prostriata tract in three individual subjects: LGN-Pro-1 (yellow) going around the lateral ventricle, reaching the LGN from a superior direction, quite likely through a set of association fibers (the tapetum) and LGN-Pro-2 (cyan) which follows a route similar to that of the OR connecting LGN with V1. On average the extracted LGN-Pro-1 comprises 23% of the fibers in the complete tract, whereas LGN-Pro-2 comprises 37% of the total number of fibers connecting LGN with prostriata (pie chart of Figure 3C, additional right hemisphere images shown in Supplementary Figure 1 and 2). It is important to note that a large majority of fibers (40%; Figure 3A) travel somewhere in between these two subcomponents, indicating that LGN-Pro is a continuous white matter bundle of which we are highlighting two components using a purely anatomically definition. We note that, this extraction is somehow arbitrary and three or more subcomponents could have been extracted from this tract. Yet, the two subcomponents we describe below comprise fibers with interesting relationship between white matter and cortical function and structure.

We further examined the microstructural properties along 100 locations of LGN-Pro-1 and LGN-Pro-2 as in the previous section. Again, the FA, MD and RD differ remarkably for LGN-Pro-1 (yellow), LGN-Pro-2 (cyan) and the complete tract (green). We also estimated the length of the complete tract (84.35±12 mm; green), LGN-Pro-1 (74.41± 7 mm; yellow) and LGN-Pro-2 (69.24±19 mm; cyan). These tract profiles show pattern similar to the ones shown by segmenting LGN-Pro-1 and LGN-Pro-2 based on their cortical termination zones (compare profiles shown in Figure 3 and 4).

Defining LGN-Pro-1 and LGN-Pro-2 independently of prostriata (based on their anatomical trajectories) as performed in this section, allowed us to analyse the distribution of the fiber terminations in prostriata. We next set to characterize any relation between the two tracts and the prostriata eccentricity, myelin-sensitivity map and cortical thickness. We found that the majority of the segregated fibers follow an eccentricity gradient as shown in the cortical maps (Figure 2). We then combine the results of our tractography analyses (Figure 4) with the gradient maps (Figure 2) to establish whether there is a relation between the white matter sub-components and prostriata.

Figure 5A shows the cortical projection zones of the two extracted white matter tracts on the cortical surface of three representative subjects (See also Supplementary Figure 3 for left hemisphere). For each individual, we projected the endpoint location of the two subcomponents on top of the gradient maps of prostriata. Qualitatively, LGN-Pro-1 appears to project more anteriorly and LGN-Pro-2 appears project more posteriorly. This division is consistent with all gradients shown within prostriata (Figure 2). In sum, given that the anterior and posterior parts of prostriata process the foveal and peripheral parts of the visual field respectively, our results suggest that whereas LGN-Pro-1 seems to transfer information from the foveal visual field, LGN-Pro-2 transfers information from the periphery. This is also consistent with the higher FA values within prostriata for LGN-Pro-1 (Figure 4B), also observed in a previous report regarding the OR retinotopic segregation (Yoshimine et al. 2018).

**Figure 5A.**
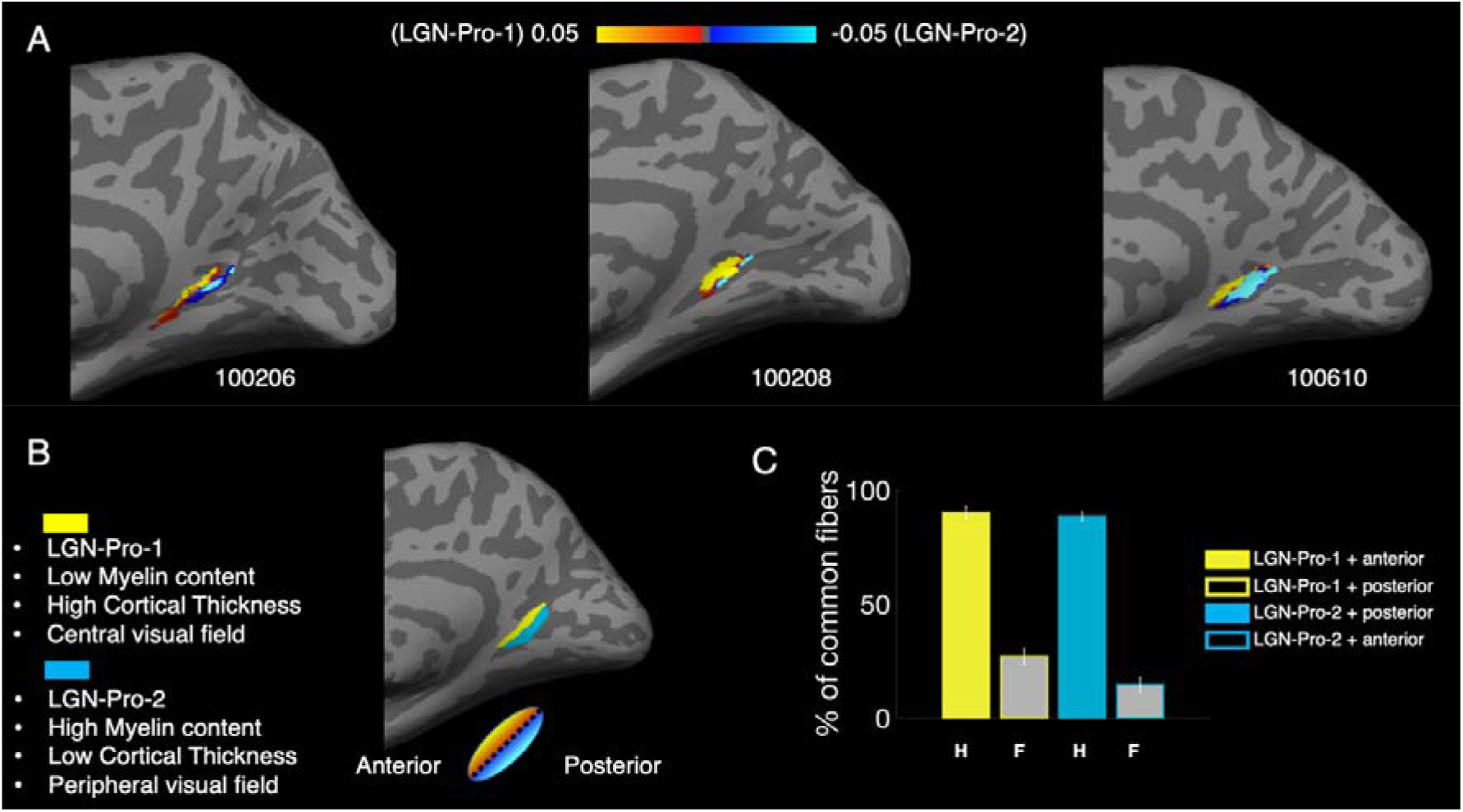
Cortical projections of LGN-Pro-1 and LGN-Pro-2 (fiber density distribution in the right hemisphere of three representative subjects). The yellow map corresponds to LGN-Pro-1 which traverses the ventricle and the blue map corresponds to the LGN-Pro-2 which followes the OR. B. Schematic summary of the LGN-Pro division based on the gradient maps. C. Overlap between LGN-Pro-1 and LGN-Pro-2. The percentage of total fibers across hemispheres separated into hits (H-colored bars) and false alarms (F-grey bars) for LGN-Pro-1 (yellow face/edge color) and LGN-Pro-2 (cyan face/edge color) which was used to calculate d’ prime.

Finally, to provide a quantification for the qualitative results reported above, we divided the prostriata ROI into two parts using the major (longer) axis of the ellipse illustrated in one representative subject in Figure 5B (this analysis is akin to that performed on 139 subjects for Figure 3). The gradient which runs along the eccentricity, myelin and cortical thickness maps was used as a guideline to divide prostriata into anterior and posterior components (see methods for details). We then counted the number of fibers entering either the anterior or posterior part of prostriata. We count as ‘Hits’ the proportion of fibers from the whole LGN-prostriata tract (in green – Figure 4A) which end or pass through the anterior part of prostriata and are identical to those in LGN-Pro-1. The proportion of fibers from the whole LGN-prostriata tract which pass through the posterior part of prostriata and are identical with those in LGN-Pro-1, are labelled as ‘False Alarms’. We compute d-prime (d’) which is the standardized difference between the means of the hits and the false alarms. In this case d’ prime is calculated to 1.90 (hits: 90.1%, false alarms: 26.9%). Then, we consider as ‘Hits’ the proportion of fibers from the green tract which pass through the posterior part of prostriata and are identical with those in LGN-Pro-2. The proportion of fibers from the whole LGN-prostriata tract which pass through the anterior part of prostriata and are identical with those in LGN-Pro-2, are considered ‘False Alarms’. In this case, d’ prime is calculated to 2.24 (hits: 88.5%, false alarms: 14.7%). Results are displayed in the bar plot of Figure 5C and demonstrate that the two subcomponents of our tract could indeed be distinguished based on their anatomical pathway.

In summary, we conclude that the anterior part of prostriata with less myelin and high cortical thickness processes information from the central visual field and is connected with the thalamus via the dorsal white matter subcomponent of the tract (LGN-Pro-1). On the other hand, the posterior part of prostriata with increased myelin and low cortical thickness processes information from the peripheral visual field and is connected with the LGN via the ventral subcomponent of the tract (LGN-Pro-2). More interestingly a similar retinotopic organization of fibers is observed in the thalamus. A retinotopic organization within LGN has been previously reported, with a foveal – peripheral gradient running from medial to lateral direction (Arcaro et al. 2015). We observe a similar organization while examining the endpoints of LGN-Pro-1 and LGN-Pro-2 within our LGN. Three example subjects are shown in Supplementary Figure 3A where LGN-Pro-1 appears to reach more medial parts and LGN-Pro-2 reaches more lateral parts of the LGN. In Supplementary Figure 4B we show a histogram with the X coordinate of the endpoints in MNI space of each subcomponent from both hemispheres of all participants (lower values – medial, higher values – lateral).

## Discussion

The aim of this study was to explore the structural connectivity of area prostriata in the human brain with the thalamus using diffusion tractography and freely available datasets from the HCP. We observe a white matter bundle connecting area prostriata with the LGN from which two subcomponents can be extracted; one that follows a similar route to the optic radiations and another one that bends around the lateral ventricle and reaches the LGN from a superior direction. We report that the fibers comprising LGN-Pro-2 are placed more posteriorly in prostriata, compared to those comprising LGN-Pro-1 which are more anterior. When comparing our anatomical results with eccentricity maps from previous reports (Mikellidou et al. 2017b; Yu et al. 2012) we observe that the peripheral part of prostriata is connected with the LGN through LGN-Pro-2 following a similar path to the OR, whereas the central visual field representation is connected through LGN-Pro-1 which circumvents the lateral ventricle. The division of the bundle into fibers that transfer information from different parts of the visual fields is also confirmed by the analysis of endpoints within the LGN. LGN-Pro-1 reaches the most medial part of LGN, which has been shown to process the foveal part of the visual field, whereas LGN-Pro-2 innervates more lateral parts, typically processing the visual periphery. Considering the resolution of data, the size of the LGN and limitation of tractography further validation would be necessary. The results obtained with the manual (Figure 4-5) and automated method (Figure 3) are consistent and taken together they demonstrate that it is possible to subdivide the LGN-prostriata white matter tract into two subcomponents, carrying information about different parts of the visual field to different parts of prostriata and the LGN.

This is not the first report of white matter tracts segmented into different subcomponent based on functional properties using in-vivo tractography. In fact, the division of a fiber bundle based on visual field representation has been carried out by Yoshimine et al.(2018), (see their Figure 2) who separate the OR based on eccentricity with clear segregation of foveal and peripheral tracts (Yoshimine et al. 2018; Parraga et al. 2012). We note that the subdivision between LGN-Pro-1 and LGN-Pro-2 is somehow anatomically arbitrary. This is because the bundle appears as a single, contiguous fibers set (Figure 3A green). Yet, it is interesting to note that retinotopically organised components can be extracted from the LGN-prostriata white matter tract similarly to what has been demonstrated for the OR.

Interestingly recent evidence shows a white matter bundle of similar shape to LGN-Pro-1 which is encompassing the ventricle, crossing the splenium and extending to the lateral occipital lobe (Yakar et al. 2018). This white matter bundle might belong to the lateral part of the tapetum, descending to the temporal lobe near the OR and the IFOF. Although prostriata is located medially in the occipital lobe, such evidence shows that the tapetum might be able to transfer information between the occipital lobe and other structures. Original work that examined the agenesis of the corpus callosum, argued that the tapetum is a separate structure and defined it as a direct continuation of the occipital frontal bundle (Onufrowicz 1887). However, subsequent work demonstrated that the tapetum is part of the splenium (Schmahmann and Pandya 2007), it consists of callosal fibers, transfers information between hemispheres (Veltin 2003) and it has been delineated in tractography studies (Martino et al. 2013; Tax et al. 2014). Irrespective of its independence from the splenium, the tapetum could be transferring visual information to the visual cortex as shown by a diffusion study that placed a seed in the splenium of the corpus callosum (Caspers et al. 2015). Interestingly, a patient with a transient splenium lesion, near the tapetum, has been reported to have a transient visual field loss with no other neurological symptoms (Gunaydin and Ozsahin 2018).

For the second part of the analysis which was performed locally we improved the anatomical constraints for tractography to avoid false positives. As the aim was to closely follow the trajectory of LGN-Pro subcomponents, we manually corrected the white matter mask for each individual hemisphere, so that the final result is limited only within voxels fully consisting of white matter. During the manual correction, we excluded all voxels with partial volume effects that were incorrectly labelled as white matter and were in fact grey matter or cerebrospinal fluid. For the larger cohort of subjects analysed on brainlife.io, we did not carry out any manual correction of the white matter mask as our aim was to determine whether subcomponents can be extracted without extensive post-processing corrections.

To map prostriata ROI on the native surfaces in the second part of analysis we used a method developed by Fischl et al. implemented in the *mri_surf2surf* function as part of Freesurfer (Fischl et al. 1999) that is based on cortical folding patterns and does not require additional functional information (unlike MSM-all). Still, we found a great degree of overlap of the projected zones when using MSM-all (Robinson et al. 2018; Robinson et al. 2014) or *mri_surf2surf* (79.4%), demonstrating that our results are not affected by alignment procedures. This is confirmed by the similarity of our results regardless of the alignment method (MSM-all versus mri_surf2surf). The two subregions of prostriata were mapped based on the HCP 7T dataset (Benson et al. 2018; Glasser et al. 2016). The stimuli presented in the study did not exceed 8 degrees of eccentricity and therefore we assume that the posterior part of prostriata is responsible for processing the peripheral and far peripheral information based on the cortical thickness and myelin sensitivity (Abdollahi et al. 2014; Burge et al. 2016). However also in this case, it is possible that the central visual field ROI extends and overlaps the abutting area located more dorsally along the POS (Elshout et al. 2018). In fact, the increased cortical thickness and decreased myelin extend in this abutting cortex that could be sharing the same central visual field representation as it is the case between V1, V2 and V3. This could suggest a different interpretation of the two tracts, with the more dorsal tract (LGN-Pro-1) associated with these new and understudied cortical areas. Either way our results suggest that associative areas have retinotopically organized projections to and/or from the thalamus and this could become important information to interpret some neuropsychological visual deficit associated with focal lesion in the white matter.

There are of course known limitations when using methods such as tractography. The emergence of false positives from tractography is not unheard of regardless of algorithms used for the analysis due to the ambiguity of diffusion MRI data (Maier-Hein et al. 2017), especially when examining structures close to crossing fibers like the sagittal striatum and OR (Goga and Ture 2015).The main weakness of tractography however is that it characterizes the axonal pathways based on indirect evidence. The approach is substantially different from the gold standards used in tract-tracing which involve physical monosynaptic transport of dyes along the axonal projections. Tracing however is only limited to brain areas of easy access for injection and is not applicable to the human brain. There has been substantial work highlighting the potential disagreement between the two methods (Pierpaoli et al. 2001; Basser et al. 1994) even when using sophisticated methods and high quality data (Thomas et al. 2014). As result, confirming the existence of any white matter connection, invasive methods and comparative work is mandatory. Indeed, in-vivo tractography findings are often complemented with comparisons to anatomical knowledge from post-mortem methods, advancing knowledge and discovery (for a discussion about statistical methods for tractography evaluation see also (Bullock et al. 2019; Pestilli 2015, 2018; Rokem et al. 2017; Rossini et al. 2019; Takemura et al. 2016b). It has been demonstrated that, under certain conditions of scientific rigor, in-vivo tractography helps to clarify the organization of white matter anatomy otherwise left unresolved by post-mortem methods (Takemura et al. 2016b; Takemura et al. 2019; Wandell 2016; Sani et al. 2019; Yeatman et al. 2014; Puzniak et al. 2019; Takemura et al. 2017; Leong et al. 2016). Furthermore, recent examples have appeared where tractography has led the way by shedding light to undescribed anatomy with postmortem work providing independent validation (Bullock et al. 2019; Kalyvas et al. 2019).

Our results demonstrate that there is a structural connection between the LGN and area prostriata in the human brain. Whether this connection is carrying an input from the thalamus, thus transferring information to prostriata cannot be answered with tractography. The LGN-prostriata connection can equally be associated with the numerous feedback projections that arise from V1 and other visual associative areas projecting monosynaptically back to the thalamus in a retinotopic-specific manner. However, electrophysiological studies in animal models do suggest the existence also of a direct thalamic input. One evidence is the high spontaneous activity and short latencies to visual stimulation of prostriata neurons in the marmoset monkey (Yu et al. 2012). In the tree shrew a direct thalamic projection to the retrosplenial cortex and specifically area prostriata have been demonstrated anatomically with thawmont autoradiography (Conrad and Stumpf 1975). Whether also in humans a direct structural input between the LGN and prostriata, bypassing V1, exists is still to be demonstrated, but if so it could be of primary importance in the presence of V1 lesions as it could be subserving residual visual abilities (Arcaro et al. 2018; Mikellidou et al. 2017a).

The distinct pattern of white matter connections carrying information from different parts of the visual field between human area prostriata and the thalamus observed here are very similar to what have been observed in V1 (Yoshimine et al. 2018). By analogy, it is tempting to suggest that similarly to prostriata all associative visual areas have retinotopically organized tracts connecting the thalamus and the cortex, demonstrating that cortical organisation extends beyond function.

## Methods

### Data from HCP and pre-processing

Pre-processed diffusion data together with anatomical T1-weighted image (AC-PC aligned) was uploaded to brainlife.io (N = 150) or downloaded locally (N = 10) from HCP bucket (Andersson et al. 2003; Andersson and Sotiropoulos 2015; Andersson and Sotiropoulos 2016; Van Essen et al. 2013). Fiber orientation density function (FOD) was estimated using Constrained Spherical Deconvolution (CSD) model (Tournier et al. 2004) with maximum harmonic order of *L*max = 10. Both CSD and diffusion tensor were processed with mrTrix3 (Tournier et al. 2019)

### Automated pipeline

#### Reproducible neuroimaging methods for automated extraction of the LGN-prostriata white matter tracts: analysis of 150 HCP subjects on brainlife.io

Brainlife.io is a free, publicly funded, cloud-computing platform that allows for developing reproducible neuroimaging processing pipelines and sharing data (Pestilli 2018; Avesani et al. 2019). To analyse the data, we developed a series of steps that are implemented as Docker containers. The pipeline was used to processing 150 subjects from the Human Connectome Project (Van Essen et al. 2013). If any of four bundles (dorsal and ventral branches for both hemispheres) was not reconstructed in automatic analysis’, we remove these subjects (N=11). We have firstly uploaded the data on brainlife.io directly from the HCP bucket and use it to align the DWI images to the T1-weighted space. To generate the ROIs (seeds for tractography) we utilized transformation files that were already available in the HCP data (Glasser et al. 2013). LGN was aligned using transformation field obtained with FSL and prostriata was transformed using MSM-all registration and workbench commands. We than fit the tensor and estimate the DOF which leads to obtaining all the necessary files for ensemble tractography (Takemura et al. 2016a). All generated tracts have been validated with LiFE (Pestilli et al. 2014) and tract subcomponents were extracted with *dtiIntersectFibersWithRoi* (minimum distance = 2 mm). Tract profiles for each subcomponent were extracted and saved (Yeatman et al. 2012). For clarity all steps are listed in a Table 1 below and code for each app is available on github (https://github.com/brainlife). For customised parameters and to perform our step-by-step procedure please access the apps using the doi links below.

| Scope of the brainlife.io App | Weblink to the online App |
| --- | --- |
| 1. Uploading DWI data together with T1-weighted ac-pc aligned anatomy on brainlife.io directly from the HCP bucket | (no app needed) |
| 2. Aligning anatomy to DWI | https://doi.org/10.25663/bl.app.68 |
| 3. Mapping ROIs from templates | https://doi.org/10.25663/brainlife.app.288 |
| 4. Fitting the tensor | https://doi.org/10.25663/bl.app.60 |
| 5. Estimating the CSD | https://doi.org/10.25663/brainlife.app.238 |
| 6. Ensemble tractography (LGN-Pros) | https://doi.org/10.25663/brainlife.app.279 |
| 7. Validating tracts with LiFE | https://doi.org/10.25663/bl.app.104 |
| 8. Intersecting the tract with ROI (dorsal/ventral prostriata) | https://doi.org/10.25663/brainlife.app.282 |
| 9. Analysis of tract profiles | https://doi.org/10.25663/brainlife.app.185 |

### Manual pipeline

#### Definition of ROIs

Cortical ROIs were firstly projected on the average anatomy ‘fsaverage’ using the template (Glasser et al. 2016) and mapped back to the native cortical space using the *mri_surf2surf*. Volume ROIs that were used as masks for tractography were created using *mri_surf2vol* command for each subject. Subcortical ROIs were mapped back from Talaraich Deamon labels included in FSL package (Lancaster et al. 2000). In the tractography session we used the following regions: the lateral geniculate nucleus (LGN) and area prostriata. The LGN ROI was dilated with *maskfilter* function (mrTrix, npass = 2). Transformations for both cortical and subcortical ROIs were obtained from HCP.

#### Tractography

For each pair of ROIs located in the same hemisphere we estimated possible white matter tracts that represent the anatomical connections between them. Tractography was constrained by the white matter mask; a union mask of two ROIs was used as a seed. For each pair we included only the fibers that traverse through both ROIs. Seed masks were also used as endpoints, ensuring that white matter connections do not simply pass through our ROIs but actually end there. Tractography was performed using mrTrix software and the *streamtrack* command. For each set of ROIs, the algorithm discovered a maximum of 10,000 fibers with 1,000,000 trials. To improve the accuracy of our results we used ensembled tractography (Takemura et al. 2016a) which performs the same tracking procedure but changes the minimum allowed curvature to propagate from voxel to voxel. We used four different curvature thresholds c = [0.25 0.5 1 2 4] mm and merged the obtained fibers in our final fiber bundles.

#### Tractography validation and thresholding

Obtained tracts were validated with LiFE software (Caiafa and Pestilli 2017; Pestilli et al. 2014). The algorithm predicts the diffusion signal using the orientation of the fascicles present in obtained connections and compare it to the acquired MR data. The difference between the two is used to calculate prediction error. For each voxel a weight is assigned that describes how each fascicle contributes towards predicting the diffusion model, with 0 signifying maximum error and 1 no error. Fascicles with zero-weights were discarded from the analysis. This procedure was applied to all tracts.

#### Manual Extraction of Prostriata subcomponents LGN-Pro-1 and LGN-Pro-2

For each subject, we used a plane extraction method to extract two subcomponents, by visually inspecting the connections and drawing an exclusion plane (all planes available as Matlab ROIs compatible with *mrDiffusion*). We focus on extracting two clearly distinct subcomponents of the LGN-prostriata tract: a dorsal and a ventral one. For example, to extract the dorsal subcomponent (LGN-Pro-1), we drew several exclusion planes (available as ROIs) to remove all fibers that travel ventrally. Similarly, to extract LGN-Pro-2 (ventral subcomponent) several exclusion planes were drawn to remove all fibers that travel dorsally.

#### Mapping the starting points

To visualize the starting points of the two subcomponents of the tract we first transformed the tractography results to the volume using *dtiFiberendpointNifti* function (vistasoft) and followed previously described surface mapping procedures from FreeSurfer (*mri_vol2vol, mri_vol2surface*). This allowed us to locate and evaluate the exact location within the seed from which fibers originate. The normalized density of fibers was calculated by taking the number of fibers passing through a voxel and dividing by the maximum number of fibers within a voxel included in the track. The density map was masked with the prostriata ROI.

#### Dividing Prostriata into anterior and posterior ROIs

Anterior and posterior parts of prostriata were manually defined using an eccentricity gradient map (Benson et al. 2018) as well as myelin and cortical thickness gradient maps (Glasser et al. 2016) (see Figure 2A-C). An example of the division of the ROI on a native surface is shown in Figure 5B. The ROI was divided in the *fsaverage* space (50% of anterior vertices and 50% of posterior vertices) and both anterior and posterior parts were mapped to the native surfaces for the purpose of this analysis.

#### Dividing the LGN-prostriata tract based on fiber endpoints within prostriata

To determine whether anatomically defined (manual method) parts of the tract transfer information from different parts of the visual field, we count the proportion of fibers of the LGN – Pro bundle entering anterior or posterior parts prostriata (see Figure 5). We defined the fibers passing through the desired ROI using *dtiIntersectFibersWithRoi.*

#### Distribution of fibers within the thalamus

To calculate the distribution of fibers inside the LGN we transformed all tracts to the MNI space (Avants et al. 2008). We then sort all the voxels inside the LGN from medial to lateral (X coordinate in MNI) and count how many fibers from each of the subcomponents enter the voxel at this specific location using *dtiIntersectFibersWithRoi* for each voxel.

#### Group averages

To create an average map of estimated tracts we utilized MNI transformation files available in the HCP project (Van Essen et al. 2013). Each ‘tck’ tract was transformed to nifti using *tckmap* function from mrtrix3 (Tournier et al. 2019) and further transformed to the MNI space. Due to the direct mapping of each individual fiber to the volume space we smoothed the map representing each tract with a FWHM = 2mm. All tracts in the MNI space were binarized and results in Figure 3 present the mean ‘consistency’ map where the value in each voxel represents the number of subjects in which the tract is present at that specific point.

