## Supplementary material for "The visual white matter connecting human area prostriata and the thalamus is retinotopically organized"

**Electronic Supplementary Material**

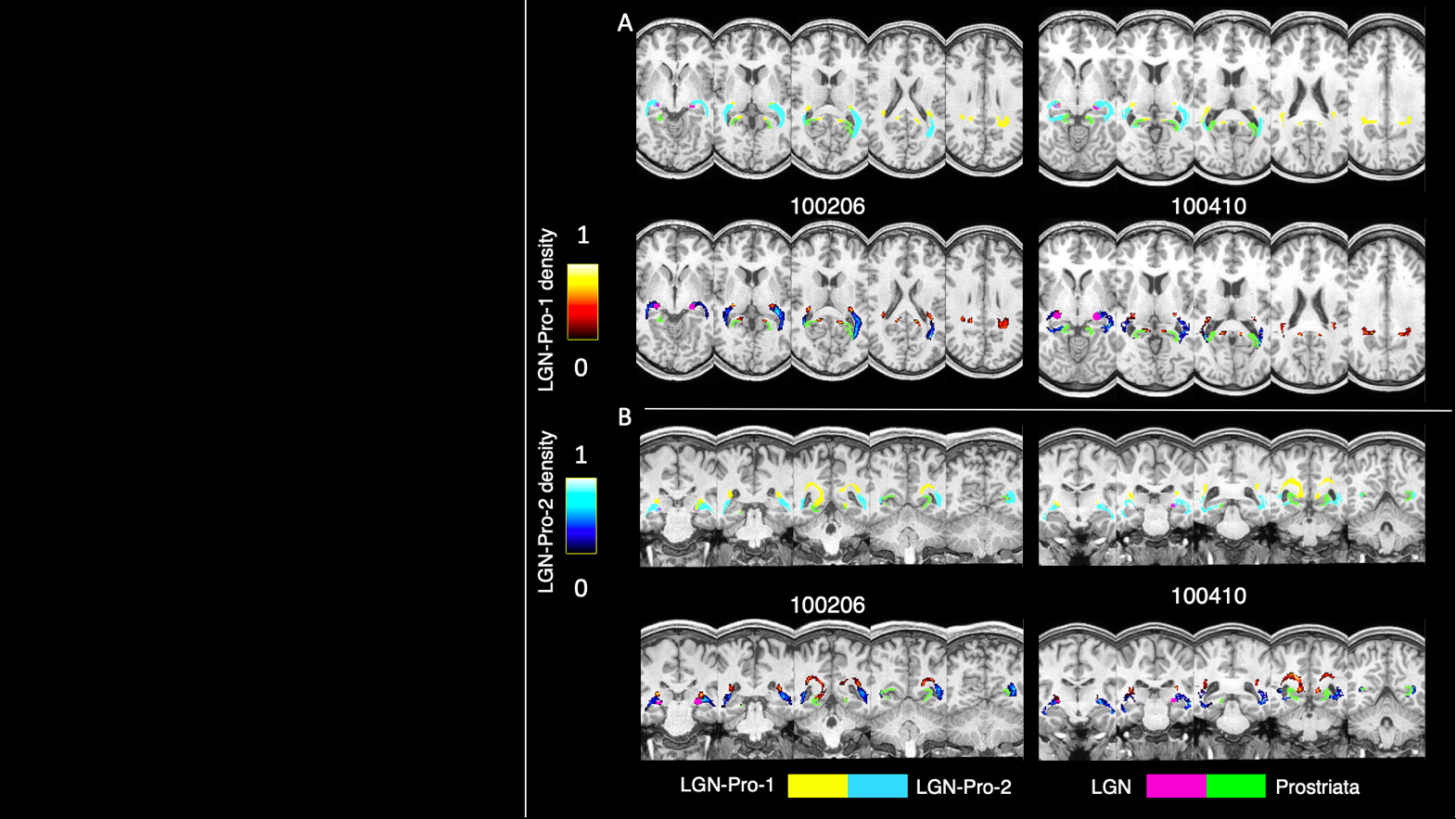

Supplementary Figure 1 - Slice by Slice tract profiles together with normalized density maps for two example subjects presented in coronal and axial planes. LGN and prostriata are outlined in pink and green colors

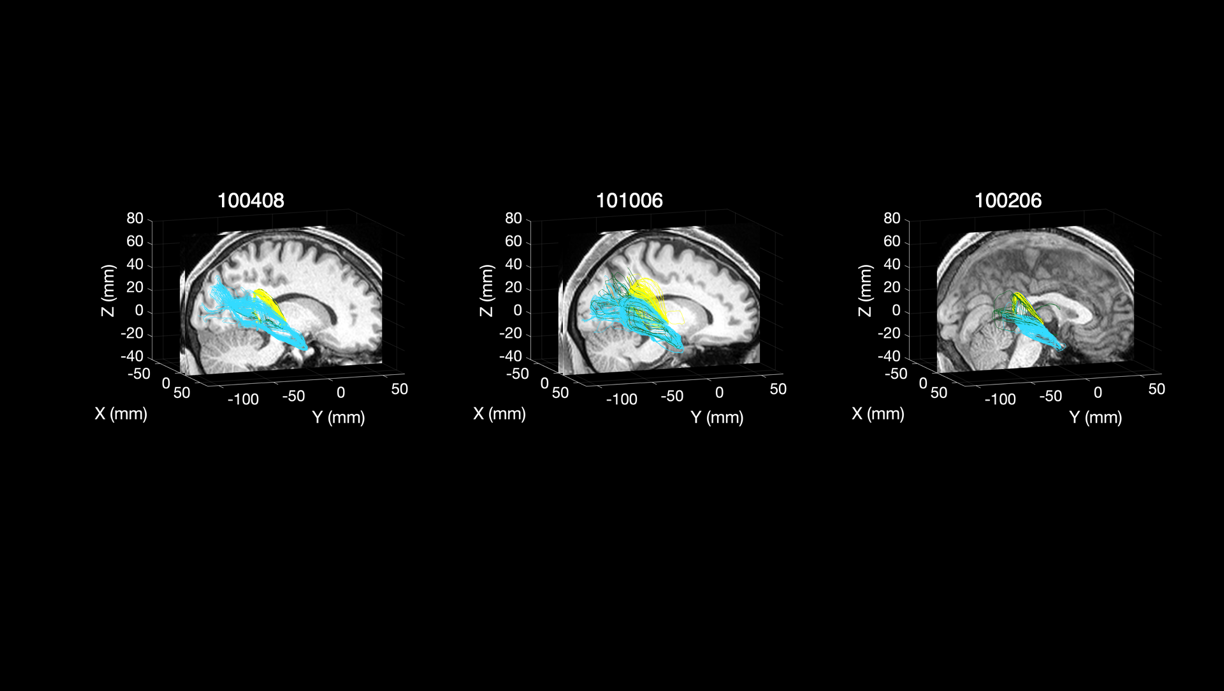

Supplementary Figure 2 – Results of the right hemispheres. Related to Figure 3A

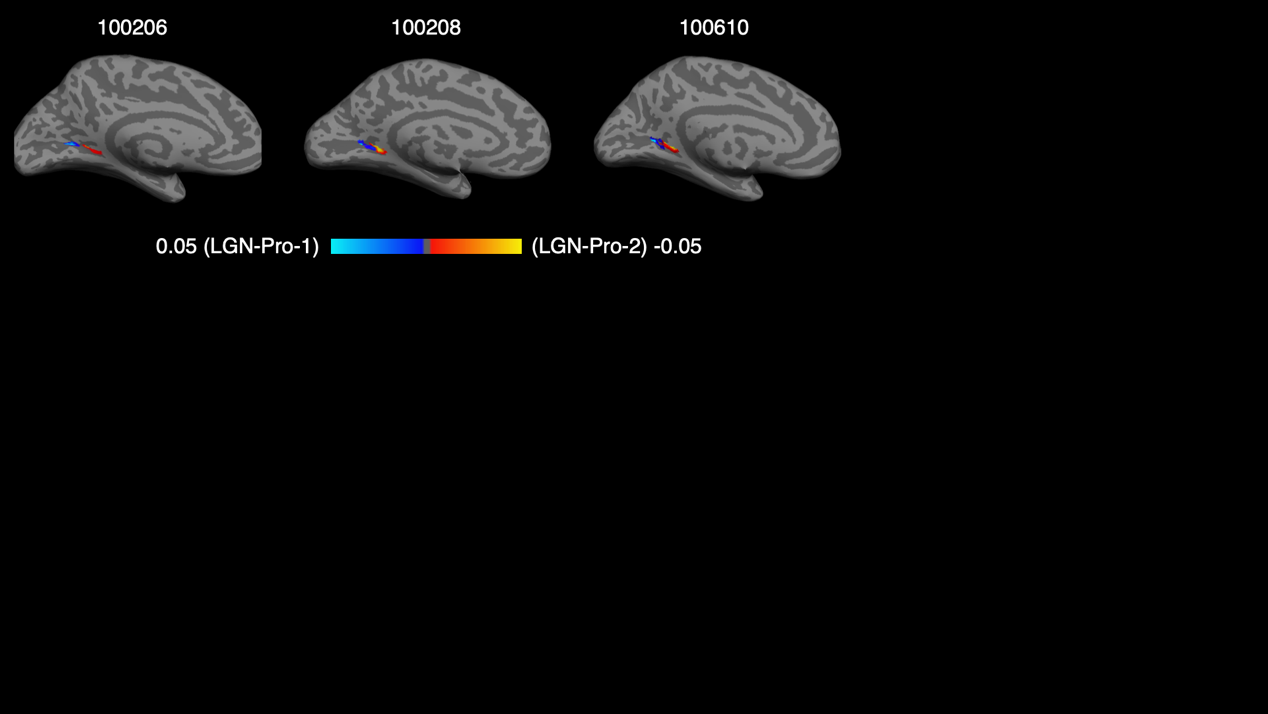

Supplementary Figure 3 – Results of the left hemispheres. Related to Figure 5A

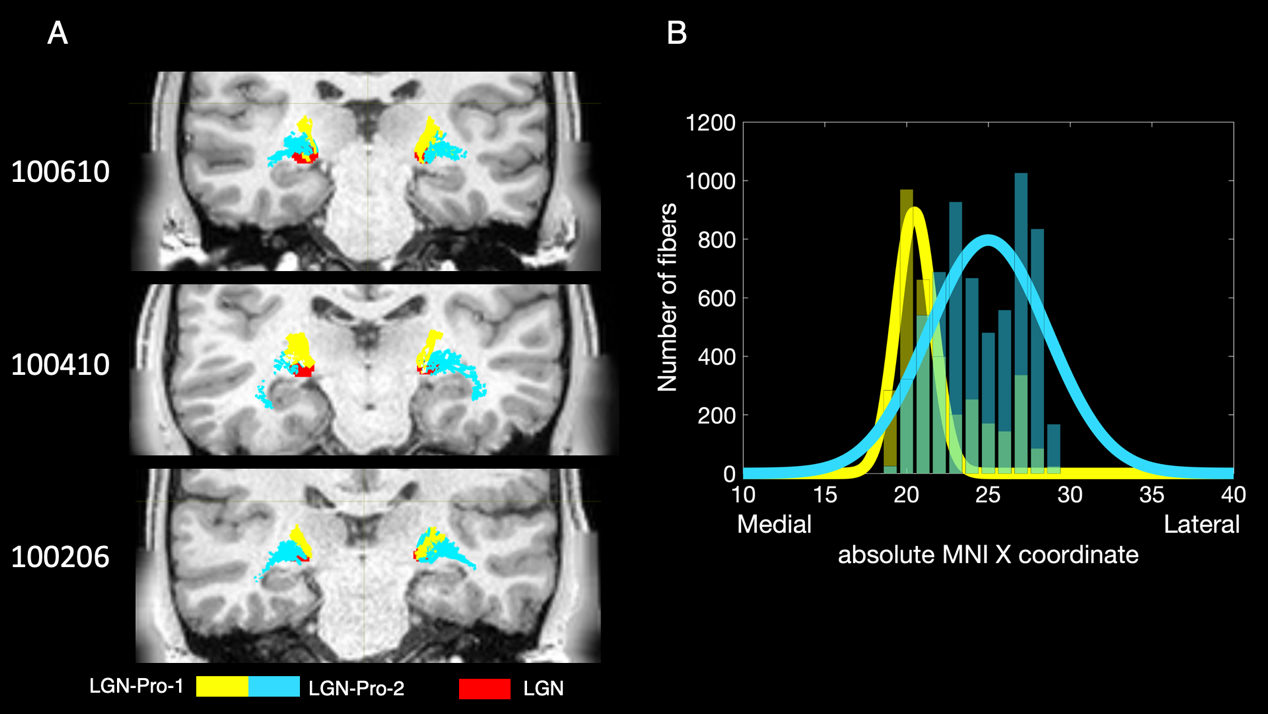

Supplementary Figure 4 - Three example subjects with a coronal slice positioned at the location of the LGN. LGN-Pro-1 enters the LGN from a superior direction reaching it medial parts while LGN-Pro-2 reaches LGN from the lateral side and finishes more laterally. B summarizes the findings as a group histogram (averaged across hemispheres) where both subcomponents are examined as a function of X coordinate in the MNI space. Thicker lines present a Gaussian fit on the binned data.
